# Transcriptional cofactor AtSDR4L and its paralog DIG2 repress somatic embryogenesis during post-embryonic development in Arabidopsis

**DOI:** 10.1101/2025.09.19.677292

**Authors:** Dongeun Go, Bailan Lu, Liang Song

## Abstract

Somatic embryogenesis, a developmental process advantageous as an alternative means for plant propagation due to the lack of maternal tissues and the ease of *in vivo* to *in vitro* knowledge transfer of embryogenesis, has been applied in studies on the regulatory mechanisms governing the developmental plasticity in plants. Core and accessory components of the polycomb repressive complexes (PRCs), the epigenetic regulatory machinery, are known to repress the transcription of genes that positively regulate the somatic embryogenesis in seedlings. Recent studies revealed that *Arabidopsis thaliana* SEED DORMANCY 4-LIKE (AtSDR4L), an interactor of a PRC accessory, represses the embryo developmental program for a successful switch from seed to seedling. However, little is known about the regulation of cell differentiation by AtSDR4L in seedlings. Here, we show that AtSDR4L, its paralog Dynamic Influencer of Gene expression 2 (DIG2), and PRC components share similar transcriptional regulatory dynamics, particularly on somatic embryogenesis-related genes. We further demonstrate that mutations in *AtSDR4L* and *DIG2* lead to the formation of somatic embryo-like structures and a delay in cellular differentiation. Thus, transcriptional changes reflected by morphological defects associated with somatic embryogenesis in *Atsdr4l dig2* mutants suggest that AtSDR4L and DIG2 prevent the dedifferentiation of cells for proper seedling growth through the transcriptional control of cell identity.

## Introduction

Plants harbour developmental plasticity, which, in part, is displayed through their ability to regenerate parts of a plant through pluripotent meristematic cells that can give rise to root and shoot or develop whole plants through totipotent embryogenic cells from somatic cells. In particular, the formation of asexual embryos arising from somatic cells, independent of fertilized gametes, occurs through somatic embryogenesis (Costa and Shaw, 2007; Fehér et al., 2016; Zimmerman, 1993). This mechanism allows clonal propagation and facilitates the study of plant embryogenesis with molecular and biochemical approaches since somatic embryo lacks maternal tissues, a feature unique in somatic embryos as opposed to zygotic embryos surrounded by and enclosed in endosperm and seed coat, respectively (Duarte-Aké et al., 2019).

Several studies in Arabidopsis have implicated that somatic embryogenesis is mediated through the transcriptional and epigenetic regulation of gene expression. For instance, overexpression of transcriptional activators such as *LEAFY COTYLEDON 1 (LEC1), LEC2, BABY BOOM (BBM)*, and *AGAMOUS-LIKE 15 (AGL15)* induces somatic embryogenesis without any additions of exogenous growth regulators (Boutilier et al., 2002; Horstman et al., 2017; Lotan et al., 1998; Stone et al., 2001; Thakare et al., 2008). Mutations in the epigenetic regulatory modules, including Polycomb Repressive Complex 2 (PRC2) components such as *CURLY LEAF (CLF)* and *SWINGER (SWN)* that function as methyltransferase for trimethylation of lysine 27 on histone H3 (H3K27me3), lead to the formation of somatic embryos (Chanvivattana et al., 2004). Likewise, mutations in ring-finger genes of the PRC1 module with monoubiquitination activity on histone H2A (H2Aub), *BMI1a/b, RING1a*/b, result in the formation of a mass of embryonic callus-like tissues (Bratzel et al., 2010; Chen et al., 2010). In addition to the somatic embryogenesis-related traits observed in mutants of *PRC* core genes, somatic embryonic calli were also observed in the mutants of *PICKLE (PKL)*, which is encoded as a chromatin remodeler, and *VIVIPAROUS/ABI3-LIKE1/2 (VAL1/2)*, which code for transcriptional corepressors, all three of which are PRC-accessories (Henderson et al., 2004; Tsukagoshi et al., 2007; Yuan et al., 2021). Disinhibition of embryonic traits and ectopic cell dedifferentiation in *prc* mutants during post-embryonic development are consistent with the derepression of embryogenesis-related genes, *LEC1, LEC2, BBM*, and *AGL15*, as well as stem cell regulatory markers, *KNAT2/6, PLT1/2*, and *WOX5/8* (Chen et al., 2010). The PRC module thus retains the differentiation of somatic cell fate by maintaining the transcriptionally silenced state of the embryonic program during postembryonic development.

*Arabidopsis thaliana* SEED DORMANCY 4-LIKE (AtSDR4L) is a transcriptional cofactor that is known to regulate embryonic and postembryonic developmental processes. *Atsdr4l* loss-of-function mutants exhibit several developmental defects, including delayed germination, seedlings with swollen hypocotyls accompanied by the overaccumulation of fatty acids in this region, as well as growth arrest with a lack of true leaves and roots (Wu et al., 2022). These pleiotropic traits overlap with those in *prc* core and accessory mutants, notably *val1 val2* (Tsukagoshi et al., 2007). De-repression of embryonic regulatory genes in *Atsdr4l* mutant seedlings, physical interaction of AtSDR4L or its paralog Dynamic Influencer of Gene expression 1 (DIG1) with VAL, and the genome-wide co-occupancy of AtSDR4L and DIG1 with VAL2 further suggest that AtSDR4L belongs to the VAL-PRC module (Lu et al., 2024; Wu et al., 2022). To date, the role of AtSDR4L as a transcriptional regulator has been examined strictly in the context of zygotic embryogenesis and the subsequent developmental events derived from this process (Cao et al., 2020; Liu et al., 2020; Lu et al., 2024; Wu et al., 2022; Zheng et al., 2022). However, our observation on the embryonic characteristics and the derepression of embryogenesis-related genes in its mutants, and the similar genome-wide binding profiles between AtSDR4L or DIG1 with VAL2, led us to speculate that AtSDR4L and its paralogs may be part of a regulatory module for the repression of somatic embryogenesis during the post-embryonic development (Lu et al., 2024; Wu et al., 2022). Here in this study, we thus reveal that AtSDR4L and its another paralog DIG2, similarly to PRC components, prevent the dedifferentiation of somatic tissues by repressing embryonic and stem cell regulatory genes for proper seedling growth.

## Results

### Transcriptome changes in *Atsdr4l dig2* seeds lay the groundwork for altered developmental path in seedlings

To examine the genes regulated by AtSDR4L in seeds and later developmental stages, we compared the *Atsdr4l-5 dig2* seed compartment transcriptome with public seedling transcriptomes of *Atsdr4l* and other epigenetic mutants that are phenotypically similar (Go et al., 2024). Clustering of the RNA-seq data shows that *Atsdr4l* seedling transcriptome is highly similar to *pkl, val1 val2, clf28 swn7, ring1a ring1b* and *bmi1a bmi1b bmi1c* (Fig. 1A). By contrast, *Atsdr4l-5 dig2* embryo and seed coat form a distinct cluster (Fig. 1A). Grouping of the differentially expressed genes by log_2_ fold changes identifies one cluster of genes with common and strong up-regulation in *Atsdr4l* single mutants and the selected *prc* mutants, while the magnitudes of de-repression are modest in the double mutant seed compartments (Fig. 1B-C). This group includes master TFs of seed maturation *ABA-INSENSITIVE3* (*ABI3*) and *FUSCA3* (*FUS3*), and a number of seed storage proteins. Intriguingly, a subset of the genes show opposite trends of misregulation between the double mutant seed compartment(s) and *Atsdr4l* seedlings, including *AGL15, NF-YC4* and a subset of the *AINTEGUMENTA-LIKE* (*AIL*) members (Fig. 1C), implying the potential stage-specific functions of AtSDR4L. Some of the genes involved in auxin biosynthesis and embryogenesis, such as *YUCCA2* (*YUC2*), *AIL5* and *BABY BOOM* (*BBM*), also displayed increased expression in both *Atsdr4l dig2* seed and *Atsdr4l* seedlings (Fig. 1D). *AIL5* and *BBM* are predominantly expressed in *Atsdr4l dig2* embryo and exhibit upregulation in both compartments. On the other hand, the master TFs of embryogenesis *LEC1, LEC2*, and *AGL15* are only significantly up-regulated in the single mutant seedlings but not in double mutant seeds, suggesting that *BBM* and *AIL5* act upstream to induce these pluripotent genes in *Atsdr4l-5 dig2*. The distribution of *ABI5* and *ABI4* transcripts differs drastically between the seed tissues and the seedlings, and *ABI5* shows weak up-regulation in both seed compartments of *Atsdr4l-5 dig2*, while *ABI4* is specifically expressed in the embryo and is not differentially expressed in the mutant (Fig. 1D). Combined, these data suggest sequential actions of AtSDR4L and its paralog DIG2 to modulate gene expression in seeds, sometimes in a tissue-specific manner, to prevent massive expression of seed maturation genes in seedlings.

**Fig. 1.**
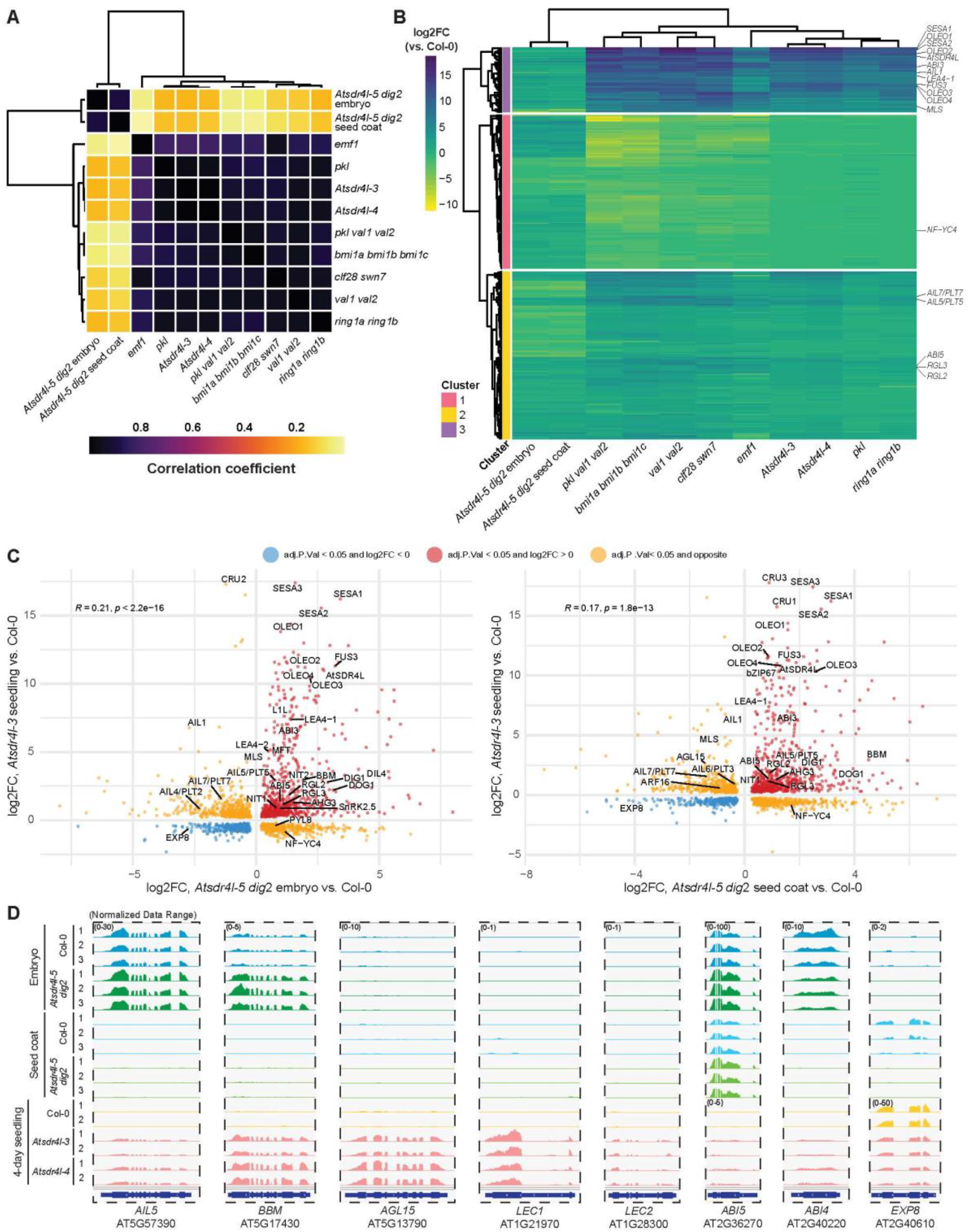
Transcriptome changes in *Atsdr4l-5 dig2* seeds lay the groundwork for altered developmental paths in seedlings. (A) Heatmap for the pairwise correlation within the transcriptomes of *Atsdr4l-5 dig2* mutant embryo and seed coat, and *Atsdr4l* and *prc* mutant seedlings. Correlation matrix was computed using log_2_FC of all shared genes among the mutants filtered by adj.P.Val < 0.05. Colour scale represents the correlation coefficients. (B) Heatmap showing the differential expression in log_2_FC (mutant vs. Col-0) of all shared significantly dysregulated genes (adj.P.Val < 0.05) among *Atsdr4l-5 dig2, Atsdr4l* and *prc* mutants. Columns representing sample genotypes and rows corresponding to genes are organized by hierarchical clustering, with labels for genes of interest related to embryogenesis, seed dormancy, storage reserve accumulation, ABA and GA metabolism and signaling, and auxin-related pathways. Colour scale reflects the log_2_FC values. (C) Scatterplot showing the distribution of log_2_FC values for significantly (adj.P.Val < 0.05) DE genes between *Atsdr4l-5 dig2* mutant embryo and *Atsdr4l-3* mutant seedling at 4 DAI (left), and between *Atsdr4l-5 dig2* mutant seed coat and *Atsdr4l-3* mutant seedling (right). Dots are color-coded for the directions of misregulation. Genes involved in seed maturation and hormone pathways of interest are highlighted. (D) Integrative Genomics Viewer (IGV) screenshots for representative genes involved in regulating embryogenesis and/or dormancy-germination switch, with similar or opposite DE trends between *Atsdr4l-5 dig2* seed compartments and *Atsdr4l* mutant seedlings. Numbers in parentheses indicate the range of normalized data in tiled data format.

**Fig. 2.**
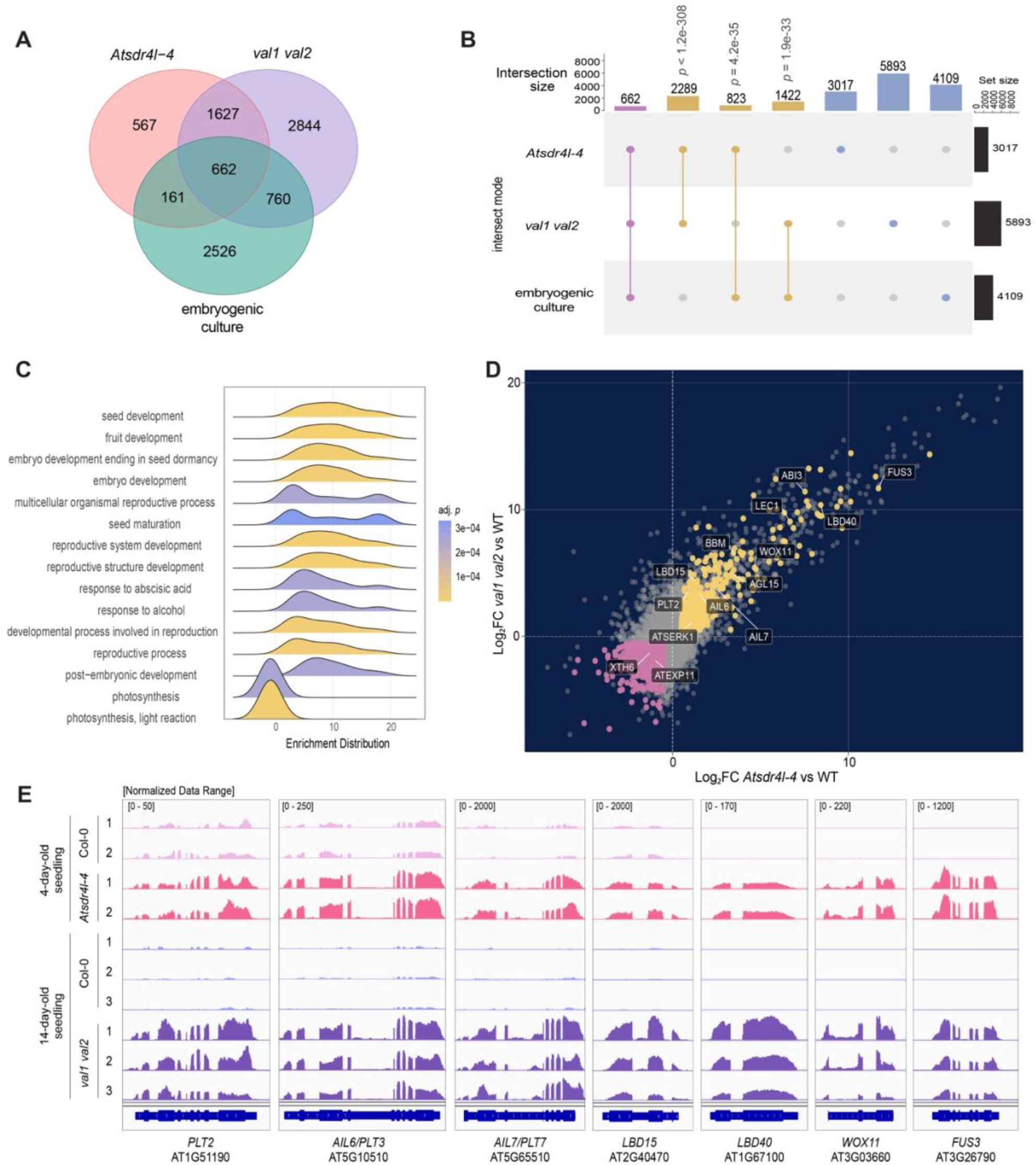
*Atsdr4l-4* seedlings, *val1 val2* seedlings, and somatic embryogenesis-induced explants with auxin harbour a shared differential expression pattern. (A) Venn diagram of significantly upregulated genes in *Atsdr4l-4* seedlings, *val1 val2* seedlings, and embryogenic cultures. (B) Upset plot of significantly upregulated genes in *Atsdr4l-4* seedlings, *val1 val2* seedlings, and embryogenic cultures. Filled circles and bars are labelled by colours as follows: the intersection among three groups (pink), the overlap between two groups (yellow), and a single set (blue). (C) Gene set enrichment plot with log_2_ fold change from the ranked list of differentially expressed genes in *Atsdr4l-4* seedlings overlapped with those in *val1 val2* seedlings and embryogenic cultures. (D) A scatter plot of log_2_ fold change from genes in *Atsdr4l-4* and *val1val2* seedlings. Each colour specifies as follows: significantly upregulated genes in *Atsdr4l-4* seedlings, *val1 val2* seedlings, and embryogenic cultures (yellow), significantly downregulated genes in all three groups (pink), and the remaining genes (grey). Selected somatic embryogenesis-related genes are highlighted in a box. (E) The read coverage of the selected genes in *Atsdr4l-4* and *val1 val2* seedlings visualized with Integrative Genomic Viewer (IGV). Numbers in brackets represent the normalized data range.

### AtSDR4L transcriptionally represses the positive regulators of somatic embryogenesis in seedlings

To date, how AtSDR4L transcriptionally repress the somatic embryonic program for proper seedling establishment has not been extensively studied. Thus, we intersected the genes that were significantly derepressed in *Atsdr4l-4* and *val1 val2* seedlings (Yuan et al., 2021) and significantly upregulated in somatic embryogenic cultures from immature zygotic embryo explants induced with auxin for 10 days (Fig. 1A,B). A significant number of depressed genes from *Atsdr4l-4* and *val1 val2* seedlings overlapped with expressed genes during the formation of somatic embryo, thereby suggesting that AtSDR4L in seedlings, similarly to VAL1 and VAL2, represses the expression of genes that are involved in somatic embryogenesis (Fig. 1A,B). The upregulated genes that were intersected in all three groups were found to be significantly enriched for embryo development and post-embryonic development, thereby further supporting that AtSDR4L in seedlings transcriptionally regulates its target genes to repress the biological processes involved in embryo formation (Fig 1C). Specifically, the transcript levels of the regulators of shoot and root identity, including *SERK1, BBM, PLT2, AIL6, AIL7, WOX11, LBD15, LBD40*, which are significantly expressed in somatic embryos, were also upregulated in both *Atsdr4l-4* and *val1 val2* seedlings (Fig. 1D,E). In alignment with the previous findings of the role of AtSDR4L-VAL2 as a repressive module of the master regulators of embryo development during seed-to-seedling transition (Lu et al., 2024), *LEC1* and *FUS3* were upregulated in all three groups as well. In contrast, cell wall loosening genes such as XTH6 and EXP11 were downregulated in all three groups, thus supporting a reduced growth of embryos in *Atsdr4l* and *val1 val2* mutants. Together, we show that AtSDR4 represses the expression of genes that positively regulate somatic embryogenesis for a successful growth of seedlings.

### AtSDR4L and DIG2 facilitate seedling establishment by promoting cell differentiation

Since *Atsdr4l and Atsdr4l dig2* seedlings exhibit multiple developmental abnormalities post-germination, further supported by a substantial number of misregulated embryogenesis-regulation genes in the mutant (Wu et al., 2022; Zheng et al., 2022), we predicted that the growth of post-embryonic development at the cellular level will also be inhibited in the mutant. Consistent with the downregulation of *LONESOME HIGHWAY (LHW)*, a TF positively regulating protoxylem differentiation, in *Atsdr4l-5 dig2* seedling (Fig. 3A), PA-PI-staining revealed that *Atsdr4l-5 dig2*, contrastingly to wild-type seedlings, lacks protoxylem at 2 DAI (Fig. 3B,C). Thus, AtSDR4L and DIG2 facilitate the differentiation of cells through the regulation of genes that control the proliferation of cells and the differentiation of vascular tissues.

**Fig. 3.**
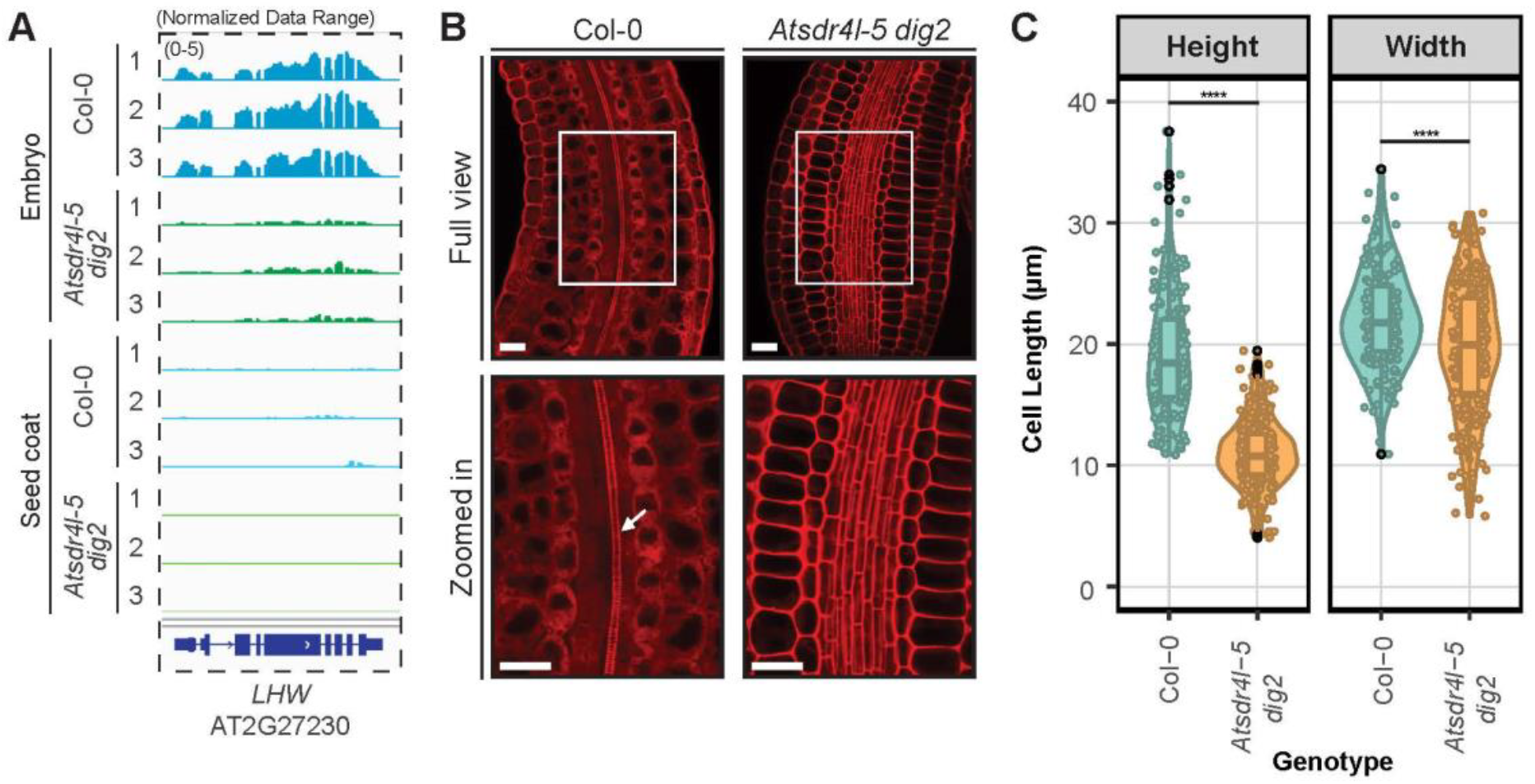
*Atsdr4l-5 dig2* embryo exhibits delayed cell differentiation and expansion. (A) Integrative Genomics Viewer (IGV) screenshots for *LONESOME HIGHWAY* (*LHW*) involved in protoxylem differentiation, with strong reduction of expression in *Atsdr4l-5 dig2* embryo. Numbers in parentheses indicate the range of normalized data in tiled data format. (B) Confocal image of periodic acid - propidium iodide (PA-PI)-stained embryos grown on 1X LS 0.7 % (w/v) phytoagar medium for 2 days. The arrow in the enlarged view of the Col-0 sample indicates protoxylem. Scale bar = 25 μm. (C) Height and width of cells at the cortex of embryos. \*\*\*\**p* < 0.0001 (48 ≤ n ≤ 89, 3 biological replicates, two-tailed Student’s t-Test).

**Fig. 4.**
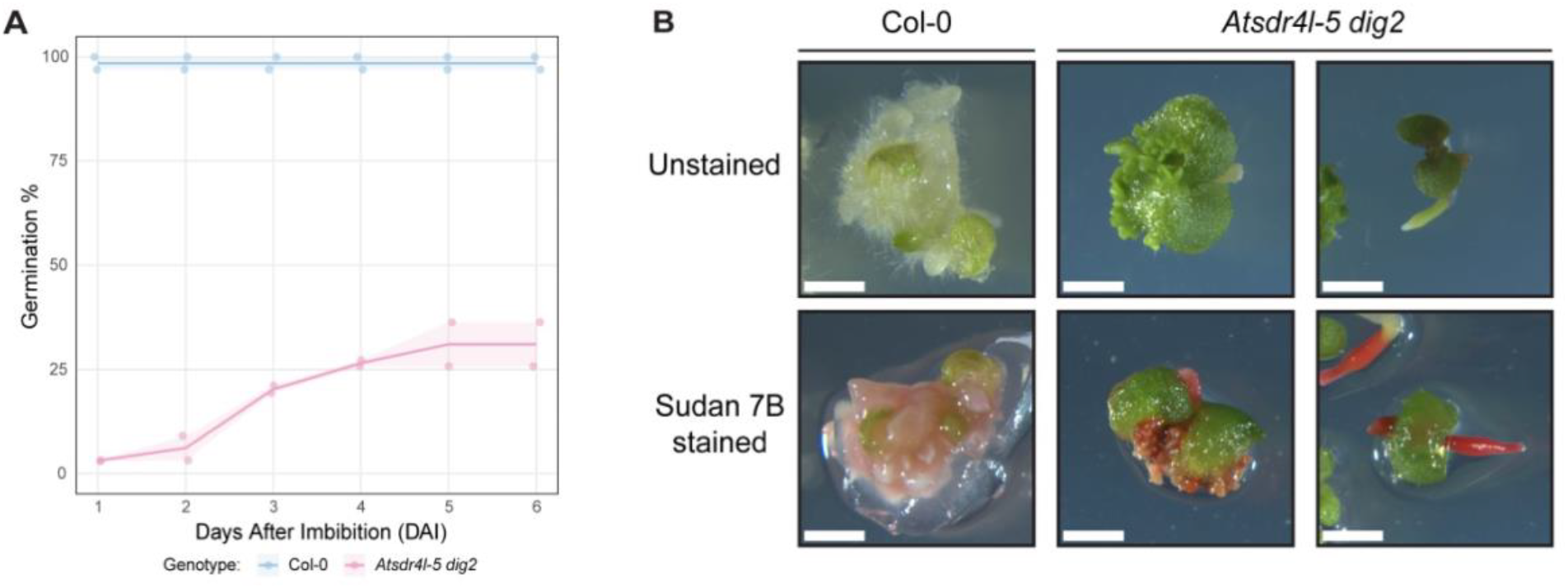
*Atsdr4l-5 dig2* embryo displays derepression of somatic embryo-like traits. (A) Germination plot of *Atsdr4l-5 dig2* and Col-0 seeds and seedlings after the transfer from auxin-containing half-strength Murashige and Skoog (MS) medium with 1% sucrose to auxin-free half-strength MS medium with 1% sucrose and 0.7 % phytoagar. Standard error of mean (31 ≤ n ≤ 33, 2 biological replicates) is represented by light background colours for each genotype. (B) A representative photo showing the growth of callus in Col-0 seedling (left). Formation of somatic-embryo-like structures from calli in *Atsdr4l-5 dig2* seedling (middle) that germinated in auxin liquid medium. *Atsdr4l-5 dig2* seedling (right) that germinated after the transfer to auxin-free medium. Scale bar = 2 mm.

### AtSDR4L and DIG2 repress the somatic embryogenesis in seedlings

The regulatory role of AtSDR4L and DIG2 in cellular differentiation and the altered transcript levels of somatic embryogenesis-regulatory genes during post-seed development prompted us to test the effect of mutation on *AtSDR4L* and *DIG2* in somatic embryogenesis during the growth of seedlings. In a hypoxic environment supplemented with auxin, a majority of *Atsdr4l-5 dig2* seeds failed to break the dormancy, while most of the Col-0 seeds successfully germinated (Fig. 1A). After a transfer to auxin-free medium, callus formation was induced in Col-0 seedlings without a subsequent formation of somatic embryos from calli whereas *Atsdr4l-5 dig2* seedlings exhibited the formation of embryo-like structures from calli (Fig. 1B). In *Atsdr4l-5 dig2* seedlings that faced minimal exposure to auxin due to the strong dormancy in culture with auxin, Sudan Red 7B stain remained in the hypocotyl and the protrusion from the shoot apex, thus indicating that the neutral lipids were over-accumulated in these regions (Fig. 1B). Hence, we show that *Atsdr4l dig2* seeds and seedlings are more responsive than Col-0 to a minimal condition for the induction of callus and somatic embryogenesis, thereby suggesting that AtSDR4L and DIG2 inhibit the formation of somatic embryos in seedlings.

## Discussion

Transcriptome analysis indicates that AtSDR4L and DIG2 help to lower the accumulation of TFs promoting embryogenesis in the embryo compartment of mature seeds. The AIL family of TFs are important regulators of cell proliferation, meristem maintenance, and embryogenesis (Horstman et al., 2014). Two *AIL* genes, *AIL2*/*BBM* and *AIL5* are commonly up-regulated between the two seed compartments of *Atsdr4l-5 dig2* and the expression level of *BBM* is considerably higher in the embryo than in the seed coat. Up-regulation of *BBM* and *AIL5* is consistent between *Atsdr4l-5 dig2* seeds and *Atsdr4l* seedlings. BBM regulates early embryo development by transcriptionally activating *LAFL* (Boutilier et al., 2002; Chen et al., 2022; Horstman et al., 2017). *AIL5* up-regulation may also be involved in the germination and embryonic phenotype, as its over-expressor is known to be hypersensitive to ABA during seed germination (Yano et al., 2009), and causes overproliferation of somatic embryos and increased transcript abundance of seed maturation genes (Tsuwamoto et al., 2010). Thus, restricting the expression of these *AIL* genes at a low level in the embryo at the end of seed maturation will favor seedling establishment over somatic embryogenesis post germination. Similarly, *FUS3* transcript abundance in mature wild-type seeds is low and the expression is largely confined to the seed coat. The up-regulation of *FUS3* and *ABI5* in the embryo and the seed coat of *Atsdr4l-2 dig2* likely explains the stronger dormancy of the mutant seeds (Alizadeh et al., 2021; Carrillo-Barral et al., 2020; Dekkers et al., 2016; Nakabayashi et al., 2012).

During seedling establishment, seed maturation genes are silenced by PRCs, and mutant seedlings of PRC core components and associated proteins exhibit a plethora of embryonic characteristics (Go et al., 2024; Ogas et al., 1997; Tsukagoshi et al., 2007; Yang et al., 2013). AtSDR4L and its paralogs may repress the expression of *LAFL* TFs through PRCs because they bind to the upstream regions of *ABI3* and *LEC1*, physically interact with VAL2, and the H3K27me3 level of *ABI3* is reduced in 3-day-old *Atsdr4l* seedlings (Lu et al., 2024; Wu et al., 2022). The link to PRCs is further strengthened by the striking transcriptomic similarity between mutant seedlings of *Atsdr4l* and PRC components. Although AtSDR4L and paralogs accumulate to high levels in nearly matured and dry seeds, PRC2 core components and VAL2 are poorly expressed at these stages (Gazzarrini and Song, 2024). This imbalance may explain the low level of H3K27me3 at *ABI3* locus and the ABA-inducibility of *ABI3* in mature and germinating seeds. By contrast, H3K27me3 level is high at *LEC1* and *LEC2* loci in mature seeds, thus these genes are difficult to activate in wild-type seedlings. In *Atsdr4l dig2* mutants, up-regulation of TFs such as *BBM, FUS*, and *ABI3* in the embryo may facilitate the reactivation of *LEC1, LEC2*, and *AGL15* in seedlings, thus changing the cell fate and developmental trajectory of mutant seedlings. Therefore, a substantial overlap and a high correlation of transcriptomic changes in *Atsdr4l dig2* mutants with those in prc core and accessory mutants, as well as the delayed cell differentiation and the formation of embryo-like structures in *Atsdr4l dig2* mutants, together suggest that AtSDR4L and its paralog DIG2, similarly to VAL of PRC module, repress the somatic embryogenesis for a successful post-embryonic development.

## Materials and Methods

### Plant materials, growth conditions, and induction of somatic embryo-like structures

Col-0 wildtype and the *Atsdr4l-5 dig2* seeds were surface sterilized and plated on 1X Linsmaier & Skoog (LS) medium (LSP03, Caisson Labs, Smithfield, UT, USA) containing 0.7% (w/v) plant agar (A111, PhytoTech Labs, Lenexa, KS, USA). Seeds were cold stratified for 3 days at 4°C and grown on plates for 2 to 4 weeks, then transferred to soil and grown under the “long-day” condition (16 h light at 23°C /8 h dark at 18°C). Yellow-to-brown siliques were harvested and stored for 2 days prior to sterilization. For induction of callus and somatic embryo-like structures, the modified protocol from Chen et al. was used (Chen et al., 2021). Sterilized seeds were inoculated in half-strength Murashige and Skoog (MS) medium (M0404, Sigma Aldrich) with 1% sucrose and 5 µM 2,4 D (D70724, Sigma Aldrich) and incubated at 4°C in the dark for 2 days. The seeds were shaken under light for 7 days at 130 rpm and were subjected to 2,4 D-free half-strength Murashige and Skoog (MS) medium with 1% sucrose and 0.7 % phytogar for two weeks. The seedlings were then stained with Sudan Red 7B for 40 minutes and underwent two rounds of wash with water, followed by five rounds of 70 % ethanol wash.

### Periodic acid-propidium iodide (PA-PI) staining

PA-PI staining of Col-0 seedlings and *Atsdr4l-5 dig2* embryos was conducted following procedures described by Truernit et al. and Sliwinska et al. (Sliwinska et al., 2009; Truernit et al., 2008) with minor modifications. After 20 weeks of after-ripening, dissected embryos, transferred on nylon mesh and grown on 1xLinsmaier & Skoog (LS) (LSP03, Caisson Labs, Smithfield, UT, USA) with 0.7 % (w/v) phytoagar medium for two days, were collected and subjected to fixative consisting of 50 % methanol (A412-1, Fisher Scientific, Waltham, MA, USA) and 10 % acetic acid (AA36289AY, Fisher Scientific, Waltham, MA, USA) at 4 C for at least 16 hours. After the wash with water, samples were incubated in 80 % ethanol. Col-0 seedlings were then incubated at 80°C for 8 minutes and *Atsdr4l-5 dig2* embryos were incubated at the same temperature for 3 minutes. Tissues were then rinsed again with water followed by incubation in fixative for 90 minutes. These samples were rewashed with water and then incubated in periodic acid (P7875-2, Sigma-Aldrich, St. Louis, MO, USA) at room temperature for 40 minutes, followed by another round of wash with water. Col-0 seedlings and *Atsdr4l-5 dig2* embryos were then incubated in Schiff reagents with propidium iodide, consisting of 100 mM sodium metabisulphite (P7875-2, Sigma-Aldrich, St. Louis, MO, USA) and 0.15 N HCI with propidium iodide (P1304MP, Invitrogen, Carlsbad, CA, USA) added to a final concentration of 100 ug/mL for 2 hours and 90 minutes, respectively. Col-0 seedlings and *Atsdr4l-5 dig2* embryos underwent the final wash with water hand incubated in a chloral hydrate solution that contains 4 g chloral hydrate (23100, Sigma-Aldrich, St. Louis, MO, USA), 1 mL glycerol (BP2290-1, Fisher Scientific, Waltham, MA, USA), and 2 mL water for 4 days and 16 hours, respectively. Finally, the samples were transferred onto microscope slides and mounted on chloral hydrate for confocal imaging.

### Image Acquisition and Processing

Confocal images of PA-PI stained Col-0 seedlings and *Atsdr4l-5 dig2* embryos were obtained using Leica sp5 confocal microscope. Tissues were visualized *via* the excitation wavelength at 514 nm and the emission wavelengths from 570 nm to 620 nm using a 40x objective under Argon laser. Images of the whole embryos from PA-PI staining, and whole seeds and embryos for germination assays and seed-coat bedding assay were acquired using Leica M205 FCA Fluorescence stereo microscope with ColorGainValueR/G/B at 8/1/21 and saturation at 25. PA-PI stained embryos were visualized with objective lens/numerical aperture (NA) at 1x/0.10 and magnification at 2.54. Seedlings used for determining the germination rate were imaged with objective lens/numerical aperture (NA) at 1x/0.02 and magnification at 0.39. Images of the embryos grown on mannitol and ABA media were captured with the objective lens/numerical aperture (NA) at 1x/0.04 and magnification at 0.82. Samples from the seed-coat bedding assay were seen with objective lens/numerical aperture (NA) at 1x/0.03 and magnification at 0.64. Acquired images were processed with Fiji software v1.54k.

### RNA-seq analysis

The public RNA-seq data for mutants of AtSDR4L and PRC-related genes were obtained from the following GEO or BioProject accessions: PRJNA663767 for *Atsdr4l-3* and *Atsdr4l-4* (Wu et al., 2022). GSE119715 for *val1 val2* (Yuan et al., 2021), GSE186152 for *pkl* (Liang et al., 2022), GSE186156 for *pkl val1 val*2 (Liang et al., 2022), GSE89356 for *bmi1a bmi1b bmi1c* (Zhou et al., 2017), and GSE154697 for *clf28 swn7, emf1-2*, and *ring1a ring1b* (Yin et al., 2021). Processing, mapping, and differential expression analyses of the public RNA-seq data were performed following the same pipeline and using the same programs as described in the bioRxiv manuscript (“Co-repressor AtSDR4L and paralog regulate hormonal and hypoxia responses in multiple Arabidopsis seed compartments”). Genes with log_2_ fold changes (LFC) > 0.5 and < -0.5 and adj.P.Val < 0.05 were considered significantly differentially expressed (DE). DEGs shared among all the mutants were extracted and intersected using tidyverse v1.3.2 (Wickham et al., 2019). Correlation matrices among the mutants were computed using the LFC values for genes with adj.P.Val < 0.05, and heatmap was created using pheatmap v1.0.12 (Kolde, 2010) with virdis colour palettes (Garnier, 2015). The LFC values were plotted by ggplot2 v3.5.1 for scatterplot (Wickham, 2011), and pheatmap v1.0.12 using the parameter cutree_rows = 3 for hierarchical clustering (Kolde, 2010) with virdis colour palettes (Garnier, 2015).

The public transcriptomic data of embryogenic cultures from immature zygotic embryo explants induced with auxin for 10 days were extracted from Wojcikowska et al. (Wójcikowska et al., 2024). Intersected genes in 4-day-old *Atsdr4l-4* seedlings, 14-day-old *val1 val2* seedlings, and embryogenic cultures of explants with auxin for 10 days were identified using VennDiagram v1.7.3 (Chen and Boutros, 2011). Upset plot of genes from these three groups was produced using the ComplexHeatmap v1.25.2 (Gu et al., 2016) by generating a combination matrix, composed of counted elements in each intersection of groups. To perform the gene set enrichment analysis, a ranked gene list from *Atsdr4l-4* seedlings that intersected with those in *val1 val2* seedlings and embryogenic cultures in decreasing order was searched against biological processes with the following parameter ‘*nPermSimple = 100000’* by ‘gseGO’ function in clusterProfiler v4.17.0 (Wu et al., 2021). Distribution of enriched genes across the rank metric was then visualized using ‘ridgeplot’ function from enrichplot v1.29.2 (Guangchuang Yu, 2018). A scatterplot displaying the log_2_ fold changes of genes in *Atsdr4l-4* and *val1 val2* seedlings was generated with ggplot2 v3.5.2 (Wickham, 2011). The visualization of read coverage tracks for each gene was performed with Integrative Genomics Browser (IGV) v2.19.5 (Robinson et al., 2011).

## Acknowledgement

The authors thank Rosado, Li, Zhang, Todesco, and Wasteneys labs in University of British Columbia for providing access to materials and equipment. The authors also thank funding support from the University of British Columbia Four Year Doctoral Fellowships (UBC 4YF) and NSERC Postgraduate Scholarship - Doctoral to B.L., UBC 4YF to D.G., and NSERC RGPIN-2019-05039 and CFI JELF/BCKDF 38187 to L.S.

